# Deletion of *Sod1* in motor neurons exacerbates age-related changes in axons and NMJs associated with premature muscle atrophy in aging mice

**DOI:** 10.1101/2022.01.27.477840

**Authors:** N Pollock, PC Macpherson, CA Staunton, K Hemmings, CS Davis, ED Owen, A Vasilaki, H Van Remmen, A Richardson, A McArdle, SV Brooks, MJ Jackson

## Abstract

Whole body knock out of Cu, Zn superoxide dismutase1 (Sod1KO) results in accelerated, age-related loss of muscle mass and function associated with a breakdown of neuromuscular junctions (NMJ) similar to sarcopenia. In order to determine whether altered redox in motor neurons is integral to this phenotype, an inducible neuron specific deletion of *Sod1* (i-mnSod1KO) was compared with wild type (WT) mice of different ages (adult, mid-age and old) and whole body Sod1KO mice. Nerve oxidative damage, motor neuron numbers and structural changes to neurons and NMJ were examined.

Deletion of neuronal Sod1 (induced by tamoxifen injection at 6 months of age) caused the exaggerated, age-associated loss of muscle mass and force generation previously reported. No effect of age or lack of neuronal *Sod1* was seen on oxidation in the sciatic nerve assessed by electron paramagnetic resonance of the *in vivo* spin probe 1-hydroxy-3-carboxy-2,2,5,5 tetramethylpyrrolidine (CPH), analysis of protein 3-nitrotyrosines or carbonyl content. i-mnSod1KO mice showed increased numbers of denervated NMJs, a reduced number of large axons and increased number of small axons compared with age-matched old WT mice. A large proportion of the remaining innervated NMJs in i-mnSod1KO mice also displayed a much simpler structure than that seen in WT mice.

Thus, while Sod1KO mice recapitulate substantially the neuromuscular phenotypes of old WT mice, deletion of Sod1 specifically in neurons induces exaggerated loss of muscle mass and force only in old (24-29 month) mice indicating that significant muscle declines require the accumulation of age-related changes such that a threshold is reached past which maintenance of structure and function is not possible.

**Significance statement:** Sarcopenia is the age-related loss of muscle mass and function. It is a significant contributor to frailty and to increased falls in the elderly. While multifactorial, changes in redox status have been shown to have significant influence over neuromuscular aging, recent work suggests that changes in motor neurons may be the driving factor in muscle atrophy. The current study confirmed that a specific lack of *Sod1* in the motor neuron causes significant alteration in axonal architecture and the neuromuscular junctions which can drive reduced muscle mass and function. Pinpointing early changes in motor neurons may provide therapeutic targets critical for maintaining muscle in the elderly.

## Introduction

The cross-sectional area of skeletal muscle in humans is reduced by 25-30% and muscle strength by 30-40% by age 70 (Porter et al., 1995). The reduction in muscle mass and function with age is due to a decrease in the number of muscle fibers and atrophy and weakening of those remaining (Lexell et al., 1986; Brooks and Faulkner, 1988; Lexell et al., 1988). The loss of muscle that occurs with aging occurs in parallel with loss of motor units in both humans and rodents (Campbell et al., 1973; Sheth et al., 2018) and is associated with a 25–50% reduction in the number of motor neurons (Tomlinson and Irving, 1977; Rowan et al., 2012; Piekarz et al., 2020). This appears to be due to selective loss of large fast α-motor neurons leading to an apparent increase in the proportion of type I (slow twitch) or type IIa muscle fibers that is particularly apparent in humans (Narici and Maffulli, 2010; Sonjak et al., 2019). Loss of innervation of individual fibers occurs in muscles with aging and we previously reported that ∼15 % of individual muscle fibers in old mice are completely denervated and ∼80 % of NMJs showed some disruption (Vasilaki et al., 2016).

In order to examine the potential role of oxidative stress and redox changes in aging, we previously examined the effect of a lack of superoxide dismutase1 (Sod1) in whole body knock out mice (Sod1KO) and demonstrated an accelerated loss of muscle mass and function that is associated with a breakdown of neuromuscular junctions (NMJ) (Jang et al., 2010; Larkin et al., 2011). Adult mice lacking Sod1 exhibit many features observed in old wild type (WT) mice (>24 months) including loss of force, altered mitochondrial function and an accumulation of structural alterations at NMJs. Subsequent studies examined the effects of specific deletion of Sod1 in muscle (mSod1KO), (Sakellariou et al., 2018) or motor neurons (nSod1KO) (Sataranatarajan et al., 2015), but both models exhibited only a mild muscle phenotype, while expression of human SOD1 specifically in neurons of Sod1KO model was found to “rescue” the phenotype such that no premature loss of muscle mass nor alterations in NMJ structure were observed (Sakellariou et al., 2014)

To address the possibility that embryonic neuronal Sod1 knockdown in the nSod1KO mice induced a compensatory effect, we have recently created an inducible motor neuron Sod1 KO (i-mnSod1KO) mouse model (Bhaskaran et al., 2020) to examine whether loss of Sod1 in motor neurons in adult-life affected age associated muscle wasting and weakness. These mice were found to have an age-related, accelerated loss of muscle mass which preceded that seen in old WT mice by 6-8 months (Bhaskaran et al., 2020).

In the current study, we examined the effects of age and lack of Sod1 only in neurons on motor neuron number and changes in nerve oxidation status as well as structural changes in axons and NMJs. Our aim was to define whether a neuron-specific lack of CuZnSOD would be sufficient to compromise neuronal oxidation, induce disruptions of neuronal and NMJ structure, and increase denervation.

## Material and Methods

### Mice

The neuron specific inducible Sod1 knock out mice (i-mnSod1KO) used in this study were generated at the Oklahoma Medical Research Foundation (OMRF) as reported (Bhaskaran et al., 2020). In order to understand the specific role of neuronal Sod1 in the premature loss of neuronal and muscle structure and mass, the i-mnSod1KO mice were compared with WT mice of 3 ages 6-9 months (adult), 16-18 months (mid-age) and 24-29 months (old). The i-mnSod1KO mice were also compared with whole body knock out (Sod1KO) mice, Sod1KO mice in which neuronal Sod1 was rescued (nerve rescue mice; SynTgSod1KO) at 6-9 months of age. Whole body Sod1KO mice and nerve rescue (SynTgSod1KO) mice were bred at OMRF as previously described (Sakellariou et al., 2014). Mice were maintained under specific pathogen-free (SPF) conditions and shipped to the University of Liverpool or the University of Michigan, where they were maintained until required. Whole body Sod1KO mice were also crossed with Thy1-CFP mice (Jackson Laboratory) at the University of Liverpool in order to produce mice (Sod1KO-Thy1-CFP) expressing CFP in neurons to allow ready visualization of the nerve without the need for antibody staining. These mice had the phenotypic changes observed in founder Sod1KO mice. The SlickH Cre mouse used to generate i-mnSod1KO mice were originally developed as a model to delete genes in motor neurons, while also simultaneously labelling the neurons with YFP and hence all expressed YFP in motor neurons (Bhaskaran et al., 2020). Male and female mice were used throughout this study.

All mice were fed ad libitum on a standard laboratory diet, subjected to a 12-h light/dark cycle and maintained under SPF conditions. All experiments were performed in accordance with UK Home Office guidelines under the UK Animals (Scientific Procedures) Act 1986 and received ethical approval from the University of Liverpool Animal Welfare Ethical Review Body (AWERB).

### Functional data

EDL force data was gathered *in situ* as previously described (Brooks and Faulkner, 1988). Briefly, with the mouse under terminal anesthesia the whole EDL muscle was isolated, the distal tendon was severed and secured to the lever arm of a servomotor and the muscle was activated by stimulation of the peroneal nerve. Maximum isometric tetanic force (Po) was recorded and specific Po was calculated from CSA. Mice were subsequently sacrificed and sciatic nerves rapidly excised and used for biochemical analyses.

### Oxidation markers

#### EPR

In order to define the activity of specific reactive oxygen species (ROS) in muscle and nerve an *in vivo* electron paramagnetic resonance spin probe protocol using 1-Hydroxy-3-carboxy-2,2,5,5-tetramethylpyrrolidine (CPH) (Noxygen) was used as previously described (McDonagh et al., 2016). The CPH probe was made up in PBS containing DETC (5µM) and DF (25µM) fresh before each procedure. Groups of mice were anaesthetised via inhalation of isoflurane (2%) and maintained at an appropriate level in this manner via nose cone throughout this procedure. Mice received a tail vein bolus injection (80µl, 9mg/kg) followed by tail vein infusion (0.225µg/kg/min) of the probe for 2hrs. The mice were sacrificed by cervical dislocation and the sciatic nerve and gastrocnemius muscle were quickly excised, stored in Krebs buffer and frozen in liquid nitrogen. Samples were stored in liquid nitrogen in a vapour phase Dewar. In order to detect levels of the oxidised probe 3-carboxy-proxyl radical (CP), samples were placed into the finger Dewar of the Bruker e-scan benchtop EPR and scanned using the settings: microwave frequency 9.78 GHz, modulation frequency 86 kHz, modulation amplitude 6.15 G, gain 10^3^.

#### Western Blot analysis

Following sacrifice, sa ciatic nerve was quickly excised and snap frozen in liquid nitrogen. The frozen nerves were cryo-pulverised under liquid nitrogen and the resulting powder added to 40µl RIPA buffer (Merck Millipore) (10x diluted with distilled water with protease inhibitors (Roche, complete, mini, EDTA-free protease inhibitor cocktail). Samples were sonicated on ice, centrifuged 12,000g for 10mins at 4°C and the supernatant retained. Protein content was quantified by BCA assay. 20µg of total protein per sample was separated by SDS-PAGE with 4% stacking and 12% resolving gels. Electrophoresis was conducted at 120V for protein separation, followed by semi-dry transfer onto nitrocellulose membrane for 1.5hrs at 300mA (Biorad Trans-Blot® SD).

A Ponceau S (Sigma) stain was used to verify the effectiveness of the transfer. The membrane was blocked using 5% fish skin gelatin made up in Tris buffered saline with tween blocking buffer (5% FSG TBS) for 1.5hrs. The membranes were incubated with primary antibody against 3-nitrotyrosines (3-NT, Abcam) (1:1000 dilution in 5% FSG TBS-T) overnight at 4°C. Duplicate gels were run to assess protein carbonyls using a kit which derivatises proteins directly on the membrane following transblotting (Cell Biolabs). For this, proteins were transferred onto PVDF membrane, and derivatised as per manufacturer’s instructions, and subsequently incubated with primary antibody against DNP (Cell Biolab; 1:1000 dilution 5% FSG TBS-T) overnight at 4°C. All membranes were then washed with TBST, and incubated with appropriate secondary antibodies for 1hr (1:20,000 dilution in 5% FSG TBST with 0.01% SDS). Protein bands were visualised on a Licor Odyssey CLx imaging system.

### Sciatic nerve cross-sections

Sciatic nerves were dissected (from point of bifurcation at the knee up to the spinal column), placed onto a needle (to prevent curling and maintain orientation) and submerged in 10% neutral buffered formalin (NBF) overnight. Nerves were transferred into a 30% sucrose solution for cryo-protection for 5 days prior to embedding in coloured OCT (Shandon Cryochrome) using cryomolds and freezing in liquid nitrogen cooled isopentane. Transverse sections (7µm) of the nerve were cut on a cryostat (Leica CM1850). Sections were thawed, blocked in 5% goat serum for 1 hour and stained using myelin protein zero (MPZ) (1 in 50 in 5% goat serum in PBS containing 1% triton) (EPR20383, Abcam) overnight. Following incubation with alexa fluor 594nm secondary slides were mounted in hard-set mountant with Dapi (Vectorlabs) and visualised on Zeiss LSM800 confocal microscope using x40 objective.

Image analysis was conducted using the free hand tool in ImageJ software (NIH). The axon and the outer edge of the stained myelin sheath of 100 axons were drawn around to provide area of the axon, area of the sheath and the distribution of axon sizes. G-ratios were calculated for each axon as the ratio of the inner axonal diameter to the total outer diameter including myelin sheath.

### Retrograde motor neuron labelling

In separate experiments, mice were anesthetized with 3% isoflurane/oxygen and the nerve branches to medial and lateral heads of the left gastrocnemius were exposed and transected. On the contralateral side the sciatic nerve was transected at the level of the mid-thigh. Motor neurons were retrograde labelled by placing sterile Gelfoam saturated with 10% FluoroRuby (Invitrogen) over the respective nerve stumps, suturing the Gelfoam in place and closing the incision sites for recovery. Post-operative analgesia was achieved by treating mice with Carprofen (5mg/kg) once a day for 3 days. Following schedule 1 cull, the lumbar regions of spinal cords were obtained by extrusion with 5 ml of PBS 7 days after the onset of labelling. Spinal cords were fixed overnight in 10% formalin at 4°C, rinsed in PBS and processed for optical clearing according to Žygelyte et. al. (Žygelytė et al., 2016) and Erturk et al., (Ertürk et al., 2012). Labelled motor neurons were imaged using an Olympus FV1000 confocal microscope with the z-step size set at 4.5 um. Z-stacks were visualized using ImageJ and motor neurons were quantified with the use of the cell counter plug in functions while scrolling through z-stacks.

### NMJ analysis

NMJ analysis was carried out on EDL muscles with 20-60 NMJs per muscle analysed. All mice utilised for these studies expressed a fluorescent protein in the nerves to allow ready visualisation. SynTgSod1KO mice were not evaluated since data relating to the NMJs in different muscles of these mice have been previously published (Sakellariou et al., 2014).

The EDL muscles were quickly excised following schedule 1 cull, pinned at roughly resting length onto sylgard plates (Fernell, Preston, UK) and fixed in 10% NBF. Whole EDL muscles were incubated with α-bungarotoxin conjugated Alexa Fluor-647nm (1:1000) (Invitrogen, Paisley, United Kingdom) in PBS (1% Triton-x) for 30minutes. Muscles were imaged via confocal microscopy (Nikon A1, Kingston, UK). The components of the NMJs were visualised using 488nm laser to excite the YFP in the i-mnSod1KO, 405nm to excite CFP in Sod1KO-CFP and the WT litter mates. Images were collected via x10 or x60 water immersion lens (Nikon). Z-stacking (1-2µm step) allowed for 3D evaluation of the NMJs.

Using NIS-elements software (Nikon) individual NMJs were identified and the ROI tool used to obtain the area of the muscle fiber that is occupied by individual NMJs in order to evaluate NMJ spreading. All NMJs were assessed and graded according to 5 separate measures of integrity and structure; (i) overlap of the pre- and post-synaptic regions (innervated, partial or denervated) and the (ii) number of post-synaptic fragments (none – no fragmentation, less than 5, 5 to 9 or greater than 10 fragments), (iii) the number of axonal sprouts and blebs, (iv) the incidence of multi-innervated NMJs, and (v) the complexity of the NMJ. This classification ranged from “complex” – showing the typical structure seen in adult WT NMJ, to “basic” – in which the NMJ appeared as simple discs with little internal structure and resembled embryonic NMJs. Two intermediate grades between complex and basic (“good” and “fair”) were also used.

### Experimental Design and statistical analysis

For all experiments the number of mice used are indicated in the associated figure legend. All statistics were carried out using Graphpad Prism 8 software, unless stated otherwise results are presented as mean ± SD.

For NMJ analysis, all NMJs present within a field of view were assigned an ID and counted/graded. Multiple 3D images per muscle were assessed and, except for the total area, all measures were expressed as a percentage of the total number of NMJs in an image in order to correct for variation in the total number of NMJs seen in each image. For fragmentation, overlap and complexity two-way ANOVA with Tukey multiple comparisons was used, while for other measures one-way ANOVA with either Dunnett’s or Tukey multiple comparisons was used.

For western blot analyses it was only possible to load n=1 onto each SDS gel for each group and in order to compare across gels the same muscle lysate was added to each and all results were normalised to this. The data were analysed via one-way ANOVA. EPR data and force were also analysed by one-way ANOVA. Significance values * p<0.05, ** p<0.01, *** p<0.001, ****p<0.0001.

Wherever possible multiple analyses were undertaken on the nerve and muscle from from each mouse. This was not possible for mice that underwent perfusion of spin probe for EPR analysis, or for used in retrograde labelling, where separate mice were examined at the University of Michigan.

## Results

The i-mnSod1KO mice were previously reported to show an exaggerated age-associated loss of *gastrocnemius* muscle mass and function (Bhaskaran et al., 2020) and the old i-mnSod1KO mice studied here showed an equivalent exaggerated loss of absolute and specific force generation in the EDL muscle (data not shown in detail).

### Oxidation markers in nerve and muscle tissue

The concentration of the EPR adduct, CP in sciatic nerves of adult Sod1KO mice was significantly increased compared with adult WT mice (Figure 1a), but WT and i-mnSod1KO mice showed no age associated changes in the amplitude of the EPR signal. Studies of the EPR signal in the EDL muscle were also undertaken and these showed an age-related increase in the WT mice. Sod1KO mice also showed a significantly higher EPR signal than WT controls (Supplementary Figure 2). In addition, in accordance with previously published data (McDonagh et al., 2016), the sciatic nerve was also observed to have a much higher EPR signal compared with skeletal muscle (Figure 1a and Supplementary Figure 1).

**Figure 1.**
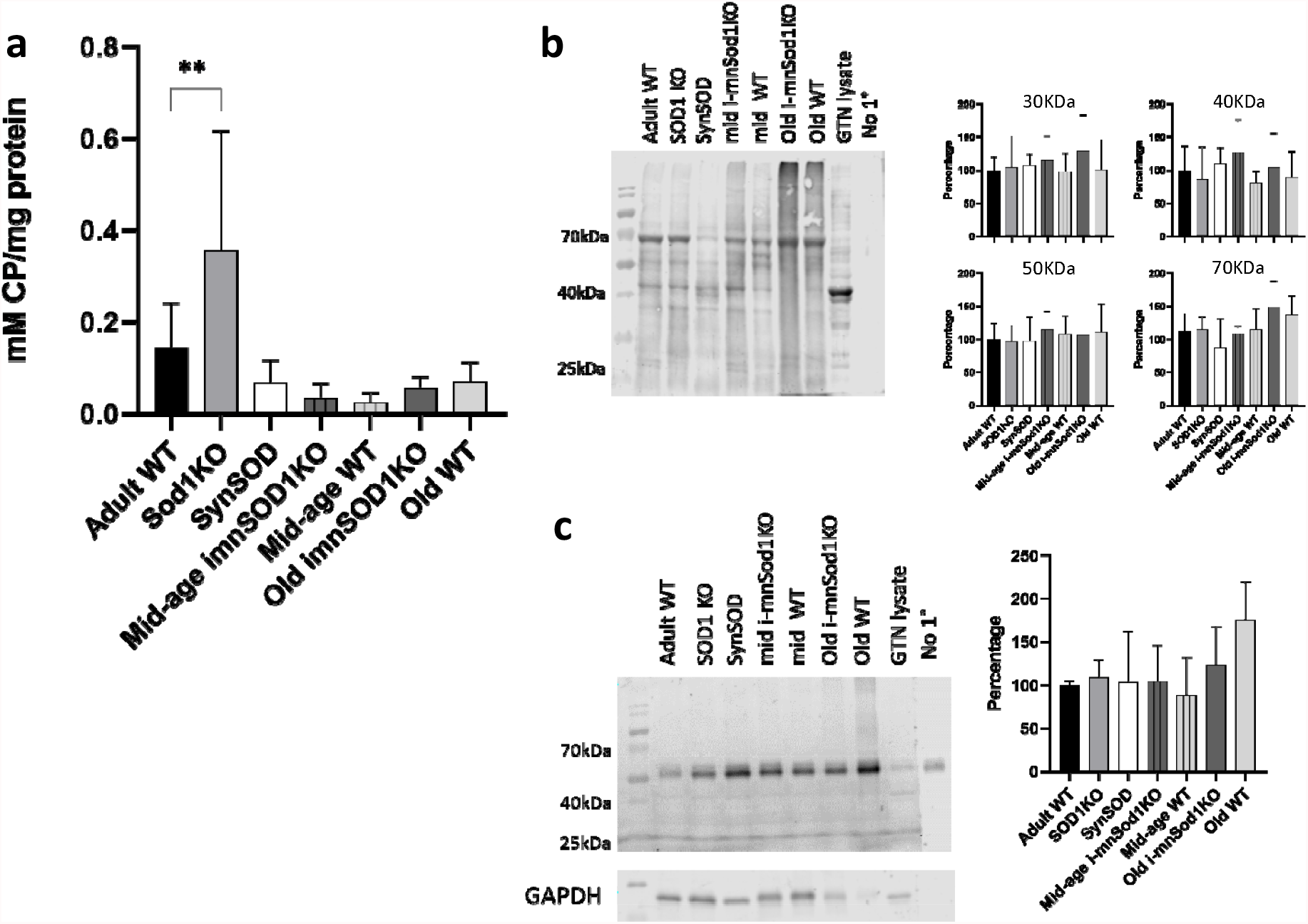
Markers of oxidation/oxidative damage in sciatic nerves of adult, mid-age and old WT, mid-age and old i-mnSod1KO, Sod1KO and SynSOD mice. Concentrations of CP, the EPR spin adduct in sciatic nerve (n=6-14) (a). Representative western blot and quantification of protein carbonyl (b) and protein 3-NT (c) contents in sciatic nerve (n=6). Data are presented as means ± SD. Symbols represent significant differences compared with adult WT ** p<0.01, from one way ANOVA analysis with Dunnett comparison. Key: GTN-lysate = lysate of mouse gastrocnemius muscle used as a positive control sample; no 1° = lane in which no primary antibody added to determine non-specific binding artefacts.

Western blot analyses for protein carbonyls (Figure 1b) revealed no statistically significant differences in sciatic nerves from any of the groups of KO or transgenic mice compared with WT mice either when the densitometry of the whole lane was assessed (data not shown in detail) or when the 4 main bands were analysed separately. Protein 3-nitro-tyrosine (3-NT) levels were also examined (Figure 1c). Few bands were detectable on the nerve western blots for 3-NT with no evidence of any change in 3NTs between the groups confirmed by densitometric analysis (Figure 1c). In these latter analyses, the major band detected (approx. 55kDa) was also present in the negative control and likely to be due to non-specific binding.

### Sciatic nerve morphology

Representative images of transverse sections through the sciatic nerve of each experimental group are shown in Figure 2a. Quantitative data for axonal size and the size of the myelin sheath around those axons was obtained. Axonal areas are shown in Figure 2b, there was no significant change in axonal area in old compared with adult WT mice, but axons from old i-mnSod1KO, Sod1KO and SynTgSod1KO mice were significantly reduced in size in comparison with age-matched WT mice. The distribution in axonal areas in each group is shown in Figure 2c and shows that there was a significant loss of the larger axons (>15µm^2^) and increase in the number of small axons in the old i-mnSod1KO, Sod1KO and SynTgSod1KO mice.

**Figure 2.**
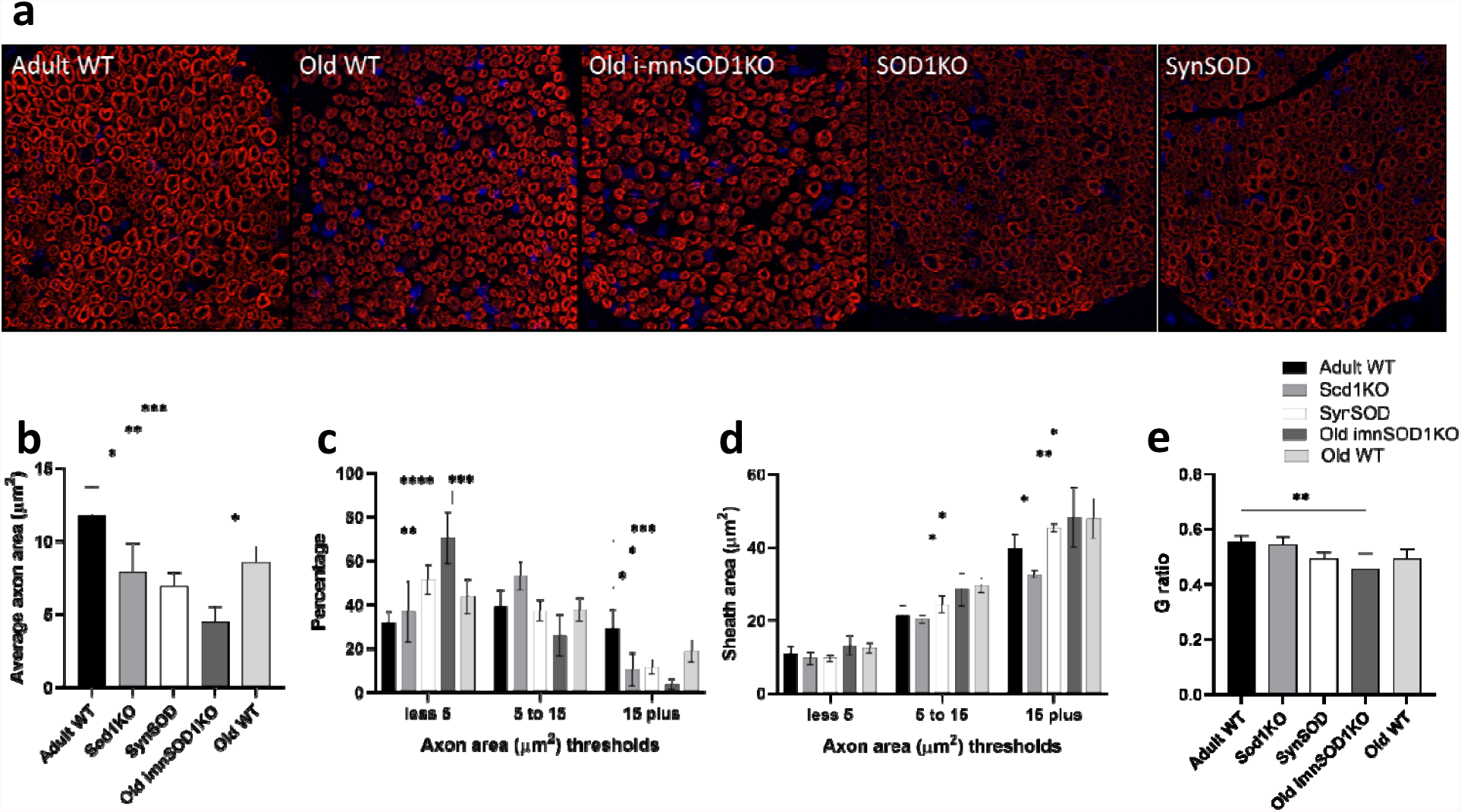
Morphology of sciatic nerves. Representative transverse cross-sections from adult and old WT mice, old i-mnSod1KO, Sod1KO and SynSOD mice are shown (a). Average axonal area (b), the distribution of axonal size (c), average myelin sheath area of axons categorised by axon size (d) and G-ratio (e) for adult, mid-age and old WT, mid-age and old i-mnSod1KO, Sod1KO and SynSOD mice. Data are presented as mean ± SD. Symbols represent significant differences (* p<0.05, ** p<0.01, *** p<0.001, ****p<0.0001) from two way ANOVA analysis with Tukeys comparison (n=4 nerves/group)

The myelin areas for different sizes of axons are presented in Figure 2d. An increase in myelin area was seen for the larger axons in the old WT mice compared with adult mice and this was also seen in the old i-mnSod1KO mice, indicating an age-related change. In contrast, there was a significant decrease in myelin area associated with the larger axons in the Sod1KO mice compared with adult WT. The G-ratio is commonly used to assess the degree of myelination and is considered a marker of efficient signal conduction (Chomiak and Hu, 2009) and when G ratios were calculated from the above data, the old i-mnSod1KO mice were found to have a significantly reduced value (Figure 2e).

### Motor neuron quantification

Images illustrating the localisation of the retrograde label to the lumbar spinal cord (Figure 3A) and to the lateral ventral horn (Figure 3B) after sciatic nerve transection demonstrate the suitability of this approach to specifically label motor neurons. Two approaches were used: labelling of the whole sciatic nerve (Figure 3C) and labelling of the nerve branches innervating the medial and lateral heads of the gastrocnemius (Figure 3D). In both situations a significant decline in the number of motor neurons was found in old compared with adult WT mice. The declines in axon numbers were also seen in old i-mnSod1KO mice. The percentage of motor neuron loss with age was approximately 30% for both gastrocnemius specific and whole sciatic nerve evaluations.

**Figure 3.**
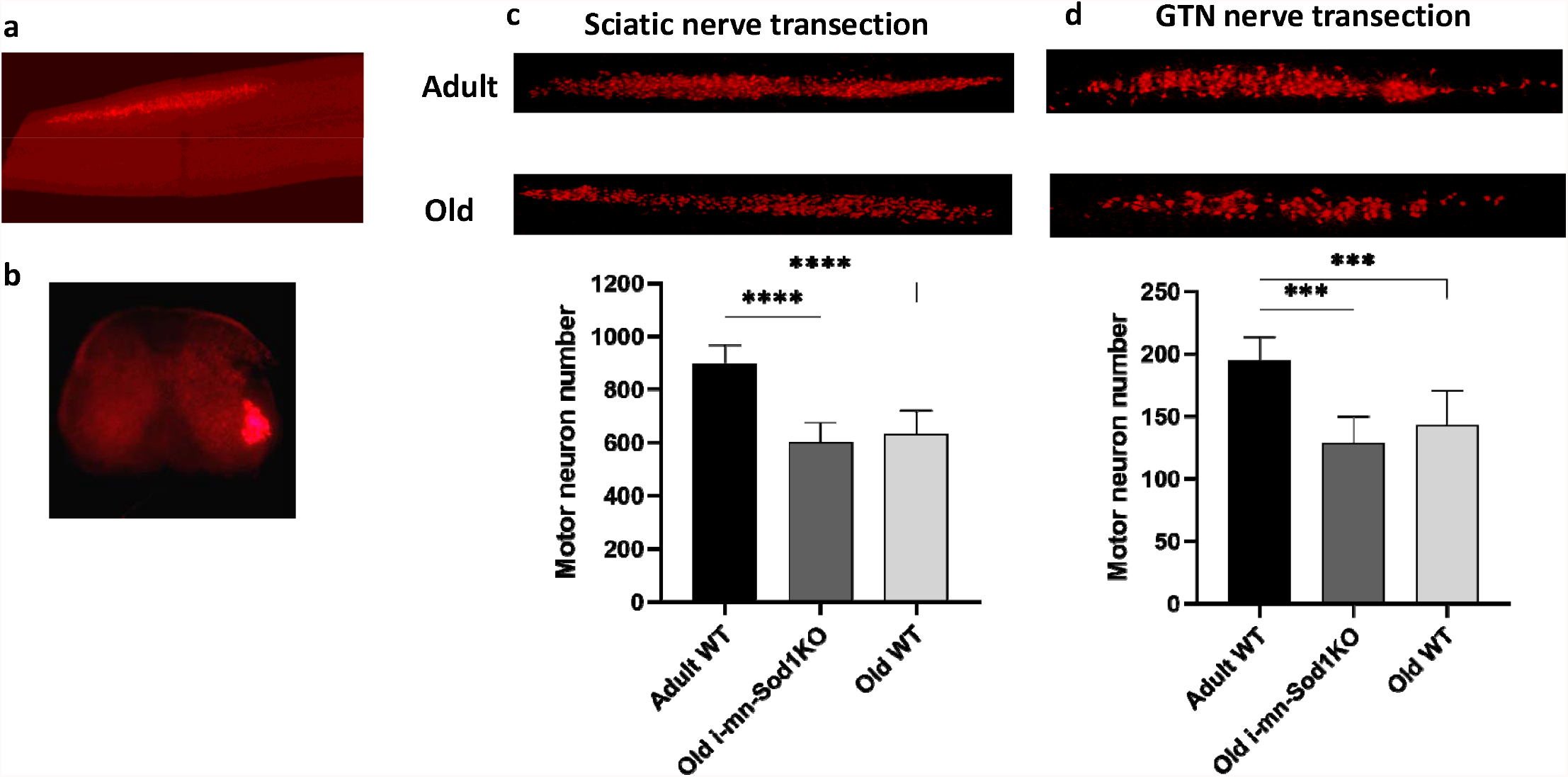
Quantification of sciatic nerve motor neurons following retrograde labelling. Representative images to show specific localisation of the retrograde label to the lumbar spinal cord (a) and to the lateral ventral horn (b) after sciatic nerve transection. Typical pattern of labelling of motor neurons in adult and old mice and quantification of numbers of labelled motor neurons following: transection of the whole sciatic nerve (c), transection of the nerve branches innervating the medial and lateral heads of the gastrocnemius (d). Data are presented as mean ± SD *** p<0.001 from 2-way ANOVA (n=3-12 animals per group).

### Changes in NMJ structure

Images of NMJs from EDL muscles were captured via confocal microscopy to allow detailed examination of their structure. Figure 4a provides representative examples from z-stacked images of NMJ from adult, mid-age and old WT mice, Sod1KO mice and mid-age and old i-mnSod1KO mice. In these images the AchR clusters were stained with bungarotoxin and the nerve visualised via CFP or YFP expression. Axonal organisation is clear and defined in adult WT but is highly disordered in both old WT and old i-mnSod1KO mice.

**Figure 4.**
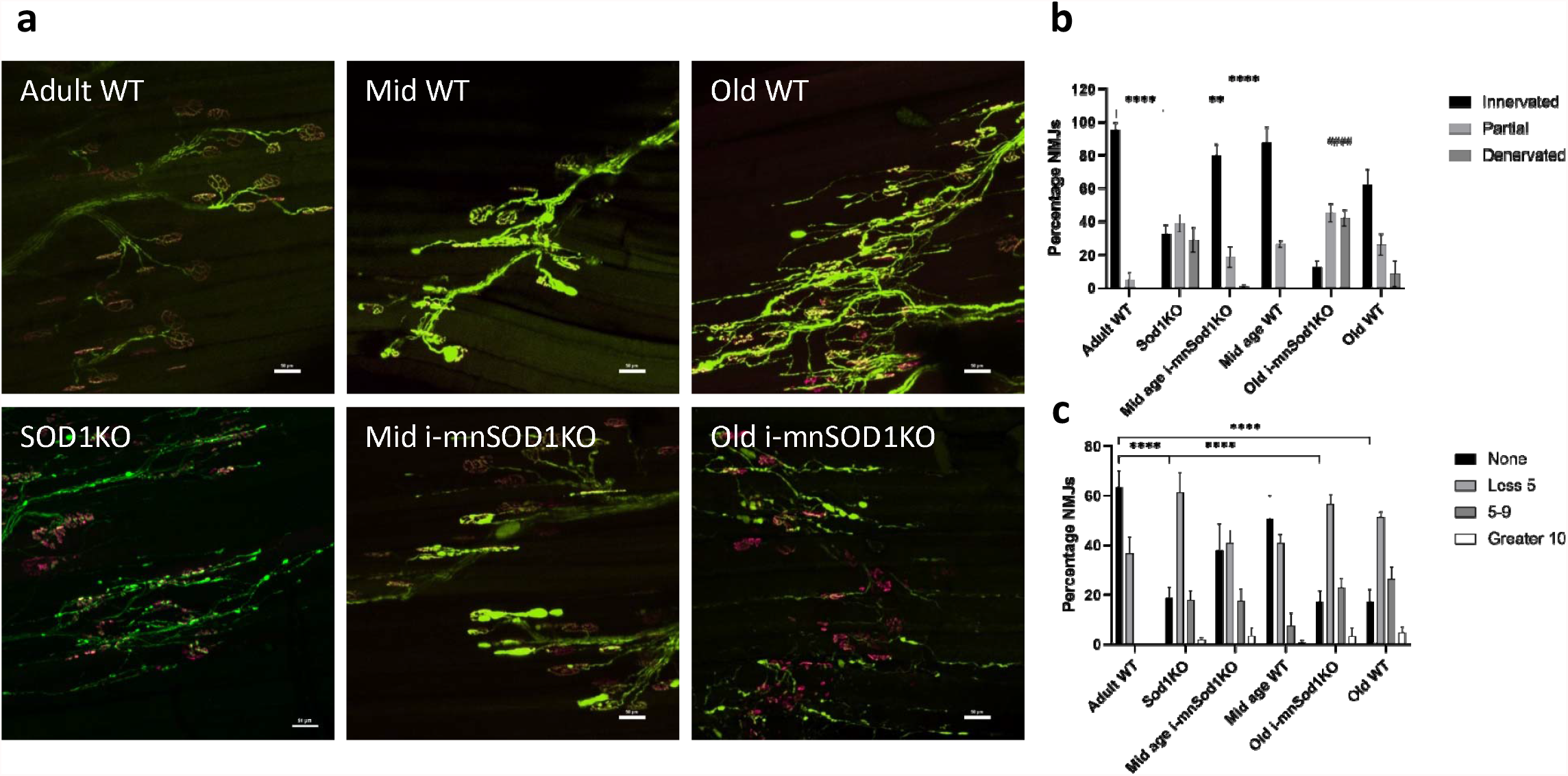
Morphological assessment of NMJ structure. Representative images of NMJ and axonal structure seen following bungarotoxin staining (magenta) and Thy1-YFP/CFP (green) visualisation in whole EDL muscles from adult, mid-age and old WT, mid-age and old i-mnSod1KO and Sod1KO-Thy1-CFP mice (a). Extent of innervation assessed by percentage overlap of pre-synaptic nerve terminal and AchR. Endplate fully innervated (black), partially innervated (light grey) or denervated (grey); (b). Average number of AchR fragments reported as none (fully intact) (black), less than 5 (light grey), 5 to 9 (grey) or more than 10 fragments (white) (c). Data are presented as means ± SD. Symbols represent significant difference compared to adult WT (** p<0.01, ****p<0.0001) or compared with old WT (^####^ p<0.0001) from two-way ANOVA with Tukey comparison (n= 4-6 muscles per group)

The average area of the muscle fiber occupied by individual NMJs showed no significant alteration either with advancing age or between any of the transgenic mouse models (data not shown). Overlap of pre- and post-synaptic regions was quantified and is shown in Figure 4b. Fully innervated NMJ had complete overlap, partially innervated NMJ meant that some of the AchRs were visible without YFP overlay and in fully denervated only AchRs were visible. The values are expressed as a percentage of the total number of NMJs counted per muscle. NMJs in muscle from adult WT mice were effectively fully innervated. NMJs from Sod1KO mice were found to be variable with roughly equivalent proportion innervated, partially innervated and denervated. At mid-age the WT and i-mnSod1KO mice had very similar profiles and showed some partially or fully denervated NMJs, but the changes were not significantly different when compared with WT adult mice. There was a significant loss of fully innervated NMJs in both old WT (p=0.0071) and old i-mnSod1KO (P<0.0001) but this loss was significantly greater in the old i-mnSod1 mice with <15% of NMJ remaining fully innervated (Figure 4b).

Fragmentation of the NMJs was quantified by counting the number of AchR clusters that were clearly separate and these were categorised as previously (Staunton et al., 2019). An intact NMJ was recorded as “none” (i.e. there were no fragments), this accounted for almost 70% of NMJs in adult WT mice (Figure 4c). The mid-aged WT and mid-aged i-mnSod1KO mice were not significantly different to each other or to adult WT mice. There was a significant decline in intact NMJs in both old groups compared with adult WT (p<0.0001) and their own mid-age groups. Sod1KO mice had only 20% of NMJ with intact AchRs, representing a significant decline compared to adult WT (p<0.0001). There were no significant differences between the age-matched groups in any of the categories.

Sprouting of axons and the presence of blebs on the axons were assessed. Figure 5a(i) shows the percentage of axons in which sprouting was evident and Figure 5a(ii) shows an example image illustrating multiple sprouting axons. There was no evidence of sprouting in the adult WT groups but the number of sprouts increased significantly in the old WT and i-mnSod1KO groups of mice. The numbers of axonal blebs in each group is shown in Figure 5b(i) and an example image illustrating multiple axonal blebs is shown in Figure 5b(ii). A significant increase in the number of neuronal blebs was seen in the Sod1KO mice and in both mid-aged WT and i-mnSod1KO mice with no significant increase in the old groups.

**Figure 5.**
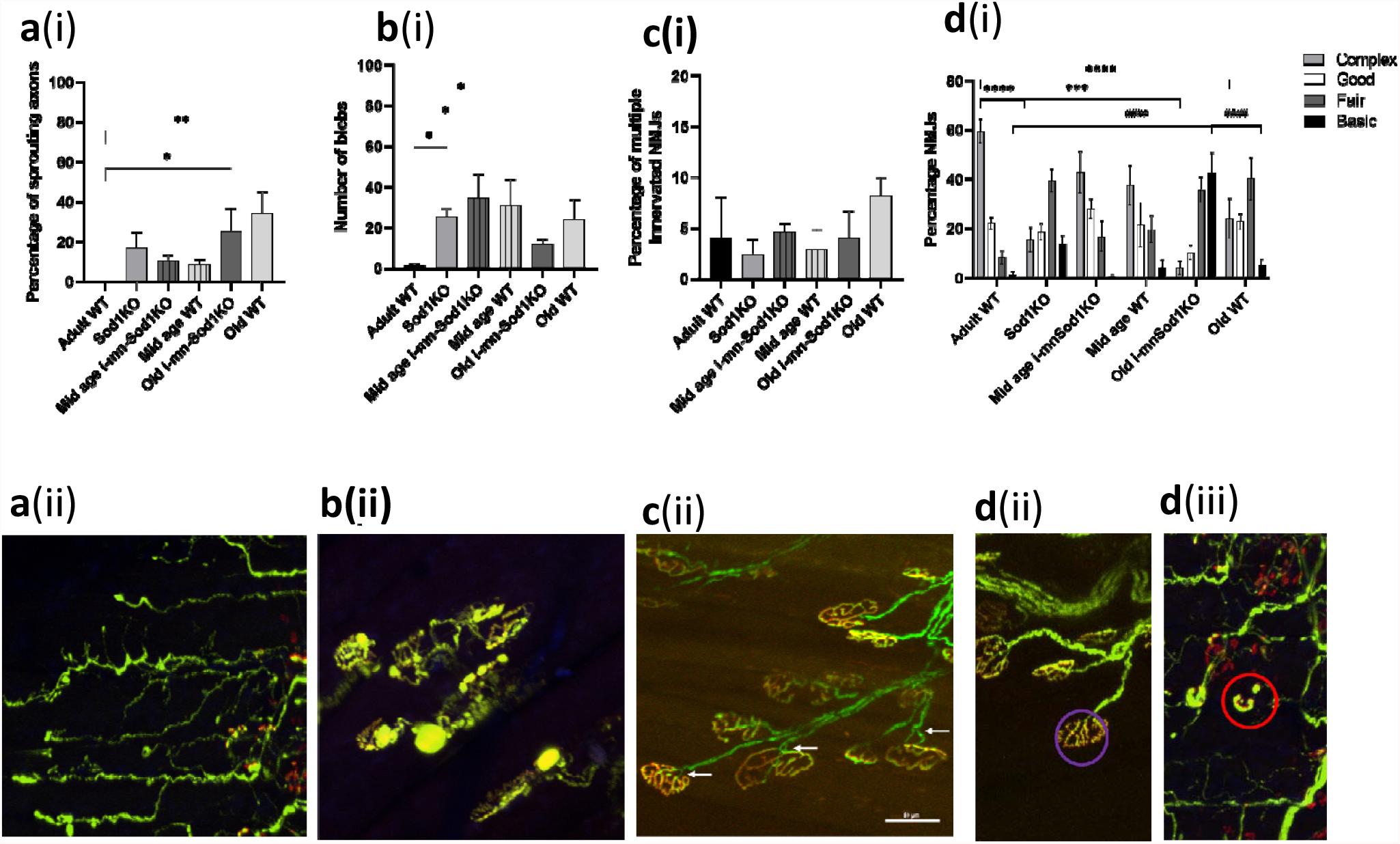
Quantification of NMJ structural alterations in EDL muscle from adult, mid-age and old WT, mid-age and old i-mnSod1KO and Sod1KO-Thy1-CFP mice. The percentage of axons having sprouts [a(i), with example of multiple sprouts at a(ii)], the number of blebs [b(i) with examples of axon with multiple blebs at b(ii)] and the percentage of NMJs which show evidence of multiple innervations [c(i) with a representative image of poly innervation at c(ii)]. Complexity of the NMJ categorised as complex (light grey), good (white), fair (grey) or basic (black); [d(i)] with a representative image of a complex [purple circle; d(ii), or basic (red circle; d(iii)] NMJ. Data are presented as means ± SD. Symbols represent significant difference in complex NMJs*** p<0.001, ****p<0.0001), or basic NMJs (####p<0.0001) from two-way ANOVA with Tukey comparison (n= 4-6 muscles per group)

Multiple innervation of single NMJs is also occasionally seen in samples from older mice and this was examined in the groups. Figure 5c(i) shows the percentage of NMJ with multiple axonal innervation and Figure 5a(ii) shows an example image illustrating these phenomena. No differences were found between the groups.

Images of NMJ show different levels of complexity with significant variability from the “pretzel shape” classically reported. A scoring system was developed to attempt to quantify and understand whether Sod1KO affected the complexity of the NMJ structure. These ranged from “complex” NMJ with numerous convolutions compared with those which were simply disks with little evidence of internal structure (Basic). Examples of complex and basic NMJ are shown in Figures 5d (ii) and (iii), respectively, and the values obtained from each group are shown in Figure 5d (i). Approximately 60% of NMJs in adult WT mice had complex junctions with less than 2% categorised as Basic (Figure 5d(i)). Old i-mnSod1KO mice had the most striking changes with approximately 40% of NMJs considered to be Basic, this was a significant increase compared with adult WT (p<0.0001) and with old WT (p=0.0003). The old i-mnSod1KO (p<0.0001) and Sod1KO (p=0.0051) mice showed a significant decrease in the percentage of complex NMJs compared with adult WT.

## Discussion

Previous studies examining the i-mnSod1KO mouse have concluded that neuron-specific deletion of CuZnSOD is sufficient to cause motor neuron loss in young mice, but loss of innervation may not be sufficient to induce muscle fiber loss until the muscle reaches a threshold beyond which it cannot compensate for neuronal loss (Bhaskaran et al., 2020). Our aim here was to define the way in which a lack of *Sod1* affects neuronal tissue to precipitate the premature age-related loss of muscle mass. Data obtained indicate that by old age, i-mnSod1KO mice showed increased numbers of fully denervated NMJ and a reduced number of large axons and increased number of small axons but no overall reduction in motor neuron numbers compared with age-matched WT mice. This was associated with changes in the morphology of the remaining innervated NMJ, which predominantly displayed a less complex structure.

A number of measures of oxidation were examined to determine whether the lack of Sod1 in nerve led to increased oxidation. We previously reported (Sakellariou et al., 2014; McDonagh et al., 2016) that 3-NT and protein carbonyls in nerves did not show a significant increase in Old WT mice, Sod1KO mice, nSod1KO or mSod1KO mice, although elevated protein carbonyls in the sciatic nerve of Sod1KO mice has been reported (Hamilton et al., 2013). This lack of any effect on nerve 3-NT or protein carbonyl contents was again seen here, but tissues were also examined using an EPR technique which previously indicated increased oxidation of the CPH probe in tissues from old compared with adult WT mice (McDonagh et al., 2016). The probe reacts preferentially with superoxide and peroxynitrite and we saw an age-related increase in muscle from WT mice (Supplementary data) but no changes in the sciatic nerve. This lack of evidence for major changes in markers of oxidative damage in nerves of i-mnSod1KO mice is compatible with our previous conclusion that the lack of *Sod1* in nerves results in impaired redox signalling, rather than oxidative damage, which plays a key role in muscle loss in Sod1KO mice (Sakellariou et al., 2018).

The previous study of i-mnSod1KO mice (Bhaskaran et al., 2020) examined motor neuron numbers in the ventral spinal cord of mice and reported early loss of motor neurons in i-mnSod1KO mice which also occurred with age in WT mice. We have used a retrograde labelling approach which is specific to individual nerves to examine motor neuron numbers. The total number of retrograde labelled sciatic nerve-associated motor neurons in adult mice is remarkably similar to that reported previously (Žygelytė et al., 2016) suggesting that it is a robust method for determining changes in motor neuron numbers. The data showed a substantial (∼30%) decline in motor neurons with age for both retrograde gastrocnemius/soleus specific and whole sciatic nerve evaluations which was not exacerbated in the i-mnSod1KO mice (Figure 3). While this technique does not differentiate between motor neuron sub-populations, alpha motor neurons make up the largest percentage of efferent neurons innervating a lower limb muscle (Burke et al., 1977; Mierzejewska-Krzyżowska et al., 2019)

Evidence that the lack of Sod1 affected axonal integrity was seen in a greater loss of axonal area in individual axons of the sciatic nerve of old i-mnSod1KO mice compared with old WT mice. These changes appeared due to a greater number of small and a reduced number of large axons in i-mnSod1KO mice in comparison with WT rather than a reduced total axon number. All older mice showed the same general trend towards reduced numbers of large and increased numbers of small axons (Figure 2).

The sciatic nerve also showed an increase in myelin thickness with advanced age as previously reported in other nerves (Peters, 2002). Both old i-mnSod1KO and age matched WT showed increased sheath thickness compared with adult mice. Neither adult Sod1KO nor SynTgSod1KO showed this accumulation of myelin. Conversely Sod1KO mice had a significant decrease in myelin sheath area in larger axons. The G-ratio (ratio of the inner axonal diameter to the total outer diameter) is a functional and structural index of optimal axonal myelination (Rushton, 1951; Chomiak and Hu, 2009) and i-mnSod1KO mice specifically showed a significantly reduced G ratio in comparison with other groups suggesting sub-optimal nerve transmission in this group. A decreased G-ratio with advancing age has been described in rats (Azcoitia et al., 2003; Amer et al., 2014)

We evaluated in detail the innervation of NMJ, fragmentation of AChRs, numbers of axonal sprouts and blebs, poly-innervation of NMJ and complexity of the NMJ structure (Figures 4 and 5). Aging caused loss of innervation, increased fragmentation, increased axonal sprouting and led to a reduced number of complex NMJs (which showed a much simpler structure). A lack of Sod1 greatly exacerbated the loss of NMJ innervation in older mice and caused an increased proportion of the NMJ to show a simpler structure, but NMJ fragmentation and axonal sprouting were unaffected by lack of Sod1. Both old WT and old i-mnSod1KO mice showed comparable changes in NMJ fragmentation, but comparison with muscle force production indicates that fragmentation may have a limited influence on signalling for force production. Furthermore, it also suggests that a lack of Sod1 in motor neurons has little influence on the post-synaptic complexes that help to maintain NMJ structure.

The lack of neuronal Sod1 in old i-mnSod1KO mice led to accelerated changes in the complexity (pretzel shape) of NMJs with age. Thus a significant percentage of NMJs in old i-mnSod1KO mice showed similar structure to embryonic (simple fragments) rather than mature pretzel shaped NMJs. It has been shown (Kang and Lichtman, 2013) that AchR clusters are lost if not re-innervated and the increased presence of NMJs with a pseudo-embryonic structure may reflect a lack of re-innervation in i-mnSod1KO mice which also showed a significant decline in NMJ innervation, in comparison with adult and old WT mice.

Thus, our data indicate that a lack of Sod1 in neuronal tissue leads to an accelerated age-related loss of skeletal muscle mass and function by accelerating or exacerbating a number of the many changes that occur in motor neurons and NMJ with aging. Although previous studies indicated that i-mnSod1KO mice showed loss of motor neurons in the ventral spinal cord of mice (Bhaskaran et al, 2000), we found no additional loss of motor neurons in excess of that which occurs in old WT mice. The assessments of motor neuron numbers in this study and that of Bhaskaran et al used different approaches, both of which are likely to have limitations that may have contributed to this divergence. For example, direct counting of motor neurons in sections of the ventral spinal cord may be limited in the ability to detect small motor neurons, while the retrograde labelling approach requires all axonal termini to be accessible to the label and the axons capable of retrograde transport. Lack of neuronal Sod1 led to a greater number of small axons and a reduced number of large axons in the sciatic nerve of old i-mnSod1KO mice in comparison with old WT and a reduced axonal G ratio suggesting impaired nerve transmission. Such changes are associated with a substantial reduction in the number of fully innervated NMJ and an increase in the number of denervated NMJ in the i-mnSod1KO mice, while some remaining AchR also show a reversion to an embryonic-type structure.

The question remains as to why the changes in nerve structure and function induced by lack of Sod1 only present as a sarcopenic phenotype in older mice, although the inducible loss of Sod1 occurs from 6 months of age. The likely explanation is that appearance of any muscle phenotype is delayed due to expansion of the size of motor units with collateral re-innervation of denervated NMJ. The time delay therefore reflects the period until the ability to expand motor unit size is exceeded. While motor units have a capacity for expansion to compensate for motor neuron loss, there appears to be a maximum ability to increase the size of an individual motor unit (Thompson and Jansen, 1977). A 25– 50% reduction in the number of motor neurons is reported to occur in both man and rodents with aging (Tomlinson and Irving, 1977; Rowan et al., 2012). In humans, Piasecki et al showed that motor unit loss occurred early in the aging process and that failure to expand was characteristic of sarcopenia (Piasecki et al., 2018). Our results are consistent motor unit loss contributing to decreased force generation with age that is exacerbated by lack of neuronal Sod1 although in both situations the initial loss of neuromuscular integrity appears mitigated by an ability to expand the size of motor units.

## Acknowledgements

The authors would like to thank the US National Institute on Aging (Grant number: AG051442), Medical Research Council (Grant number: MR/M012573/1) and the University of Liverpool for their generous financial support. Thanks also to the Biomedical Services Unit, University of Liverpool for excellent animal care throughout the project.

---

**Supplementary Figure 1.**
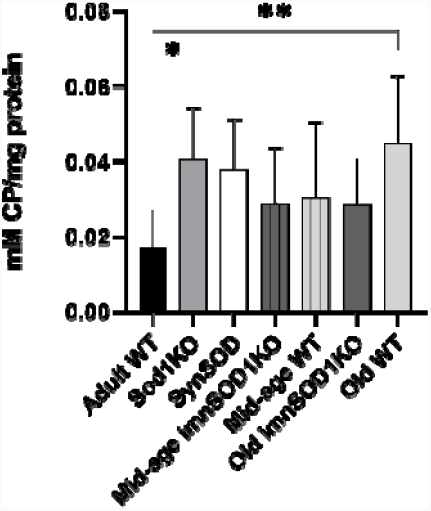
EPR analysis of the concentration of CP in skeletal muscle from adult, mid-age and old WT, mid-age and old i-mnSod1KO, Sod1KO and SynSOD mice (n=6-14). Data are presented as mean ± SD. Symbols represent significant differences (* p<0.05, ** p<0.01, *** p<0.001, ****p<0.0001) from one way ANOVA analysis with Tukeys comparison

## References

Amer MG, Mazen NF, Mohamed NM (2014) Role of calorie restriction in alleviation of age-related morphological and biochemical changes in sciatic nerve. Tissue Cell 46:497–504.

Azcoitia I, Leonelli E, Magnaghi V, Veiga S, Garcia-Segura LM, Melcangi RC (2003) Progesterone and its derivatives dihydroprogesterone and tetrahydroprogesterone reduce myelin fiber morphological abnormalities and myelin fiber loss in the sciatic nerve of aged rats. Neurobiol Aging 24:853–860.

Bhaskaran S, Pollock N P CM, Ahn B, Piekarz KM, Staunton CA, Brown JL, Qaisar R, Vasilaki A, Richardson A, McArdle A, Jackson MJ, Brooks SV, Van Remmen H (2020) Neuron-specific deletion of CuZnSOD leads to an advanced sarcopenic phenotype in older mice. Aging Cell:e13225.

Brooks SV, Faulkner JA (1988) Contractile properties of skeletal muscles from young, adult and aged mice. J Physiol 404:71–82.

Burke RE, Strick PL, Kanda K, Kim CC, Walmsley B (1977) Anatomy of medial gastrocnemius and soleus motor nuclei in cat spinal cord. J Neurophysiol 40:667–680.

Campbell MJ, McComas AJ, Petito F (1973) Physiological changes in ageing muscles. J Neurol Neurosurg Psychiatry 36:174–182.

Chomiak T, Hu B (2009) What is the optimal value of the g-ratio for myelinated fibers in the rat CNS? -DA theoretical approach. PLoS One 4:e7754.

Ertürk A, Becker K, Jährling N, Mauch CP, Hojer CD, Egen JG, Hellal F, Bradke F, Sheng M, Dodt HU (2012) Three-dimensional imaging of solvent-cleared organs using 3DISCO. Nat Protoc 7:1983–1995.

Hamilton RT, Bhattacharya A, Walsh ME, Shi Y, Wei R, Zhang Y, Rodriguez KA, Buffenstein R, Chaudhuri AR, Van Remmen H (2013) Elevated protein carbonylation, and misfolding in sciatic nerve from db/db and Sod1(-/-) mice: plausible link between oxidative stress and demyelination. PLoS One 8:e65725.

Jang YC, Lustgarten MS, Liu Y, Muller FL, Bhattacharya A, Liang H, Salmon AB, Brooks SV, Larkin L, Hayworth CR, Richardson A, Van Remmen H (2010) Increased superoxide in vivo accelerates age-associated muscle atrophy through mitochondrial dysfunction and neuromuscular junction degeneration. FASEB J 24:1376–1390.

Kang H, Lichtman JW (2013) Motor axon regeneration and muscle reinnervation in young adult and aged animals. J Neurosci 33:19480–19491.

Larkin LM, Davis CS, Sims-Robinson C, Kostrominova TY, Van Remmen H, Richardson A, Feldman EL, Brooks SV (2011) Skeletal muscle weakness due to deficiency of CuZn-superoxide dismutase is associated with loss of functional innervation. Am J Physiol Regul Integr Comp Physiol 301:R1400–1407.

Lexell J, Downham D, Sjöström M (1986) Distribution of different fibre types in human skeletal muscles. Fibre type arrangement in m. vastus lateralis from three groups of healthy men between 15 and 83 years. J Neurol Sci 72:211–222.

Lexell J, Taylor CC, Sjöström M (1988) What is the cause of the ageing atrophy? Total number, size and proportion of different fiber types studied in whole vastus lateralis muscle from 15-to 83-year-old men. J Neurol Sci 84:275–294.

McDonagh B, Scullion SM, Vasilaki A, Pollock N, McArdle A, Jackson MJ (2016) Ageing-induced changes in the redox status of peripheral motor nerves imply an effect on redox signalling rather than oxidative damage. Free Radic Biol Med 94:27–35.

Mierzejewska-Krzyżowska B, Celichowski J, Bukowska D (2019) The differences in the structure of the motor nucleus of the medial gastrocnemius muscle in male and female rats. Ann Anat 221:93–100.

Narici MV, Maffulli N (2010) Sarcopenia: characteristics, mechanisms and functional significance. Br Med Bull 95:139–159.

Peters A (2002) The effects of normal aging on myelin and nerve fibers: a review. J Neurocytol 31:581–593.

Piasecki M, Ireland A, Piasecki J, Stashuk DW, Swiecicka A, Rutter MK, Jones DA, McPhee JS (2018) Failure to expand the motor unit size to compensate for declining motor unit numbers distinguishes sarcopenic from non-sarcopenic older men. J Physiol 596:1627–1637.

Piekarz KM, Bhaskaran S, Sataranatarajan K, Street K, Premkumar P, Saunders D, Zalles M, Gulej R, Khademi S, Laurin J, Peelor R, Miller BF, Towner R, Van Remmen H (2020) Molecular changes associated with spinal cord aging. Geroscience 42:765–784.

Porter MM, Vandervoort AA, Lexell J (1995) Aging of human muscle: structure, function and adaptability. Scand J Med Sci Sports 5:129–142.

Rowan SL, Rygiel K, Purves-Smith FM, Solbak NM, Turnbull DM, Hepple RT (2012) Denervation causes fiber atrophy and myosin heavy chain co-expression in senescent skeletal muscle. PLoS One 7:e29082.

Rushton WA (1951) A theory of the effects of fibre size in medullated nerve. J Physiol 115:101–122.

Sakellariou GK, Davis CS, Shi Y, Ivannikov MV, Zhang Y, Vasilaki A, Macleod GT, Richardson A, Van Remmen H, Jackson MJ, McArdle A, Brooks SV (2014) Neuron-specific expression of CuZnSOD prevents the loss of muscle mass and function that occurs in homozygous CuZnSOD-knockout mice. FASEB J 28:1666–1681.

Sakellariou GK, McDonagh B, Porter H, Giakoumaki, II, Earl KE, Nye GA, Vasilaki A, Brooks SV, Richardson A, Van Remmen H, McArdle A, Jackson MJ (2018) Comparison of Whole Body SOD1 Knockout with Muscle-Specific SOD1 Knockout Mice Reveals a Role for Nerve Redox Signaling in Regulation of Degenerative Pathways in Skeletal Muscle. Antioxid Redox Signal 28:275–295.

Sataranatarajan K, Qaisar R, Davis C, Sakellariou GK, Vasilaki A, Zhang Y, Liu Y, Bhaskaran S, McArdle A, Jackson M, Brooks SV, Richardson A, Van Remmen H (2015) Neuron specific reduction in CuZnSOD is not sufficient to initiate a full sarcopenia phenotype. Redox Biol 5:140–148.

Sheth KA, Iyer CC, Wier CG, Crum AE, Bratasz A, Kolb SJ, Clark BC, Burghes AHM, Arnold WD (2018) Muscle strength and size are associated with motor unit connectivity in aged mice. Neurobiol Aging 67:128–136.

Sonjak V, Jacob K, Morais JA, Rivera-Zengotita M, Spendiff S, Spake C, Taivassalo T, Chevalier S, Hepple RT (2019) Fidelity of muscle fibre reinnervation modulates ageing muscle impact in elderly women. J Physiol 597:5009–5023.

Staunton CA, Owen ED, Pollock N, Vasilaki A, Barrett-Jolley R, McArdle A, Jackson MJ (2019) HyPer2 imaging reveals temporal and heterogeneous hydrogen peroxide changes in denervated and aged skeletal muscle fibers in vivo. Sci Rep 9:14461.

Thompson W, Jansen JK (1977) The extent of sprouting of remaining motor units in partly denervated immature and adult rat soleus muscle. Neuroscience 2:523–535.

Tomlinson BE, Irving D (1977) The numbers of limb motor neurons in the human lumbosacral cord throughout life. J Neurol Sci 34:213–219.

Vasilaki A, Pollock N, Giakoumaki I, Goljanek-Whysall K, Sakellariou GK, Pearson T, Kayani A, Jackson MJ, McArdle A (2016) The effect of lengthening contractions on neuromuscular junction structure in adult and old mice. Age (Dordr) 38:259–272.

Žygelytė E, Bernard ME, Tomlinson JE, Martin MJ, Terhorst A, Bradford HE, Lundquist SA, Sledziona M, Cheetham J (2016) RetroDISCO: Clearing technique to improve quantification of retrograde labeled motor neurons of intact mouse spinal cords. J Neurosci Methods 271:34–42.

